# Apical and basolateral plasma membranes in epithelial cells have distinct lipidomes and biophysical properties

**DOI:** 10.1101/2025.11.20.689573

**Authors:** Carolyn R. Shurer, Kandice R. Levental, Ilya Levental

## Abstract

Epithelial cell polarization is essential for many physiological processes, including tissue morphogenesis, nutrient absorption, barrier integrity, and directional secretion. A defining feature of such polarization is the separation of plasma membrane lipids and proteins into distinct apical and basolateral compartments. It has long been suggested that the apical compartment is rich in glycolipids and cholesterol and that this composition arises through trafficking of self-assembled membrane domains (termed lipid rafts). However, neither the detailed composition nor the mechanisms of protein and lipid sorting between plasma membrane compartments in epithelial cells have been fully resolved. Particularly, the lipid profile of the basolateral membrane, and consequently the lipid disparity between the apical and basolateral membrane, remain undefined. The general determinants of protein sorting are also poorly understood. We developed a novel method to separately isolate the apical and basolateral plasma membranes and used lipidomics and biophysical profiling to characterize the changes in membrane composition and properties between these compartments in polarized Madin-Darby canine kidney cells. We find that the apical membrane is enriched in cholesterol, saturated lipids, and glycolipids relative to the basolateral membrane, and that its biophysical properties reflect a more ordered environment. Further, we evaluate the longstanding hypothesis that lipid rafts contribute to apical protein trafficking by assessing the relationship between transmembrane domain raft affinity and apical localization. We observed that lipid raft affinity only modestly influences apical versus basolateral sorting. These findings define the distinct compositional and biophysical features of apical and basolateral compartments of epithelial cells and provide mechanistic evidence for their biogenesis.

## INTRODUCTION

Epithelial cell polarization is essential for the development and maintenance of complex tissues, enabling directional transport, selective permeability, and tissue compartmentalization^1^. The defining feature of epithelial polarization is the segregation of the plasma membrane (PM) into distinct apical and basolateral compartments, separated by tight junctions that severely restrict molecular exchange of both proteins and lipids^2^. The apical PM acts as a robust barrier protecting tissue from noxious external conditions and regulates nutrient absorption and waste excretion. The basolateral PM functionally connects epithelial cells to their neighbors and the endocrine signaling system. Consistent with these divergent functions, the protein components of the two epithelial PM compartments are highly distinct^3^, though how proteins are sorted between them remains poorly understood, with most previous investigations focusing on a few specific cases^4–8^. Clathrin has emerged as an important trafficking component in basolateral protein sorting^3,9^. However, apical sorting signals remain poorly defined, though they appear to involve glycans and other features that interact with lectins, lipid rafts, and cytoplasmic motor proteins^4,10^. Recent work has suggested a size-based sorting mechanism in which proteins with short cytoplasmic tails are preferentially sorted to the apical PM^11^, indicating an important role for biophysical mechanisms in maintaining polarized protein distributions.

Studies on epithelial cells have revealed extensive lipid remodeling during polarization, supporting the long-standing idea that apical and basolateral PM compartments possess distinct lipid compositions. Classical studies of *ex vivo* colonic tissue were among the first to note the apical PM’s unusually high abundance of sphingolipids (SLs)^12^. More recently, high-resolution mass-spectroscopy based lipidomics confirmed the relative enrichment of SLs (at the cost of glycerophospholipids (GPLs)) in the apical PM relative to whole cells^13^. Further, the lipid profile of epithelial cells was reported to evolve during polarization, with progressive changes in headgroup profiles, hydrophobic chain length, and lipid unsaturation during *in vitro* polarization^14^. While these groundbreaking studies demonstrated complex dynamics of lipid profiles during epithelial polarization, there has not yet been a detailed, quantitative comparison between the lipid compositions of the apical and basolateral PMs, leaving open the question of the lipidomic disparity between these closely associated PM compartments.

Lipid compositions dictate functional physical properties of membranes such as lipid packing, membrane fluidity, and membrane tension^15–17^. These properties contribute to cellular processes by regulating protein function, signaling, diffusion, and organization, as well as bulk physical properties like trans-bilayer permeability and membrane rigidity^18^. Although membrane properties are widely considered essential for epithelial barrier integrity and directional transport, they have not been systematically compared between distinct polarized PM compartments. While measurements with fluorescent reporters have suggested differences in lipid packing within epithelial cells^19,20^, comprehensive, quantitative comparisons between the biophysical properties of apical and basolateral membranes in fully polarized epithelial cells remain scarce.

A historically prominent hypothesis for sorting of lipids and proteins to the apical PM involves lipid rafts, tightly packed membrane domains that self-assemble due to preferential interactions between saturated SLs and cholesterol. The raft hypothesis was originally formulated as a mechanistic explanation for the unique lipid and protein composition of the apical PM in epithelial cells^12^, proposing that rafts coalesce in the trans-Golgi and therein provide a sorting platform for transport of selected (i.e. raft-preferring) proteins and lipids to the apical PM^21–24^. This hypothesis of raft-mediated sorting was later expanded to also encompass PM traffic in non-polarized cells^25–29^. However, despite the persistence and stature of this hypothesis, there has been no systematic comparisons between protein raft affinity and polarized PM sorting.

In this study, we address these gaps by developing a novel approach to specifically isolate apical and basolateral PMs and applying mass spectrometry-based lipidomics, biophysical profiling, and protein engineering to systematically compare the lipid composition, membrane properties, and protein partitioning between the PM compartments of polarized epithelial cells. We show that apical and basolateral compartments of MDCK cells differ in lipid composition, with the apical PM enriched in raft-associated lipids and exhibiting tighter lipid packing as reported by environment-sensitive dyes. We also find that proteins with higher raft affinity tend to localize more to the apical PM; however, this effect is relatively modest in comparison with raft-associated sorting to the PM in general. Together, our findings provide quantitative evidence for biophysical mechanisms involved in the organization and function of polarized epithelial PMs.

## RESULTS

### Apical and basolateral plasma membranes have distinct lipidomes

To compare apical and basolateral PM lipidomes, we isolated giant plasma membrane vesicles (GPMVs), which can be induced to bud from the PM of intact cells by chemical treatments^30^. Such GPMVs have been widely used for lipidomic analyses of PM and are highly enriched in the lipids and proteins of the PM^28,31,32^. To isolate apical and basolateral PMs, we first produced polarized epithelial cells by growing MDCK-II cells on transwell filters for ≥13 days (cell polarization was confirmed via segregation of classical apical and basolateral markers, (Supporting Fig 1a). GPMVs were induced in this setting and collected from either the apical or basolateral of the transwell filters (Fig 1a). We validated that the tight junctions that normally restrict the flow of proteins and lipids between the apical and basolateral compartments maintained their integrity during GPMV formation and isolation by selectively staining the two sides of the cell monolayer with different lipophilic dyes (e.g., red DiI on the apical side and green DiO on the basolateral side) before generating GPMVs. Flow cytometry revealed that these dyes do not mix between the two vesicle populations (Supporting Fig 1b-d). Western blotting for endoplasmic reticulum (ER) protein marker calnexin revealed minimal ER contamination (Supporting Fig 1e and Supporting Fig 2), consistent with previous validation of GPMVs as highly PM enriched preparations^33^. Finally, Western blotting for established apical and basolateral protein markers revealed little cross-contamination between the PM compartments (Supporting Fig 1e and Supporting Fig 2). These observations support the conclusion that GPMVs isolated from polarized MDCKs represent highly enriched apical and basolateral PMs, referred to hereafter as ApPM and BlPM, respectively.

**Figure 1.**
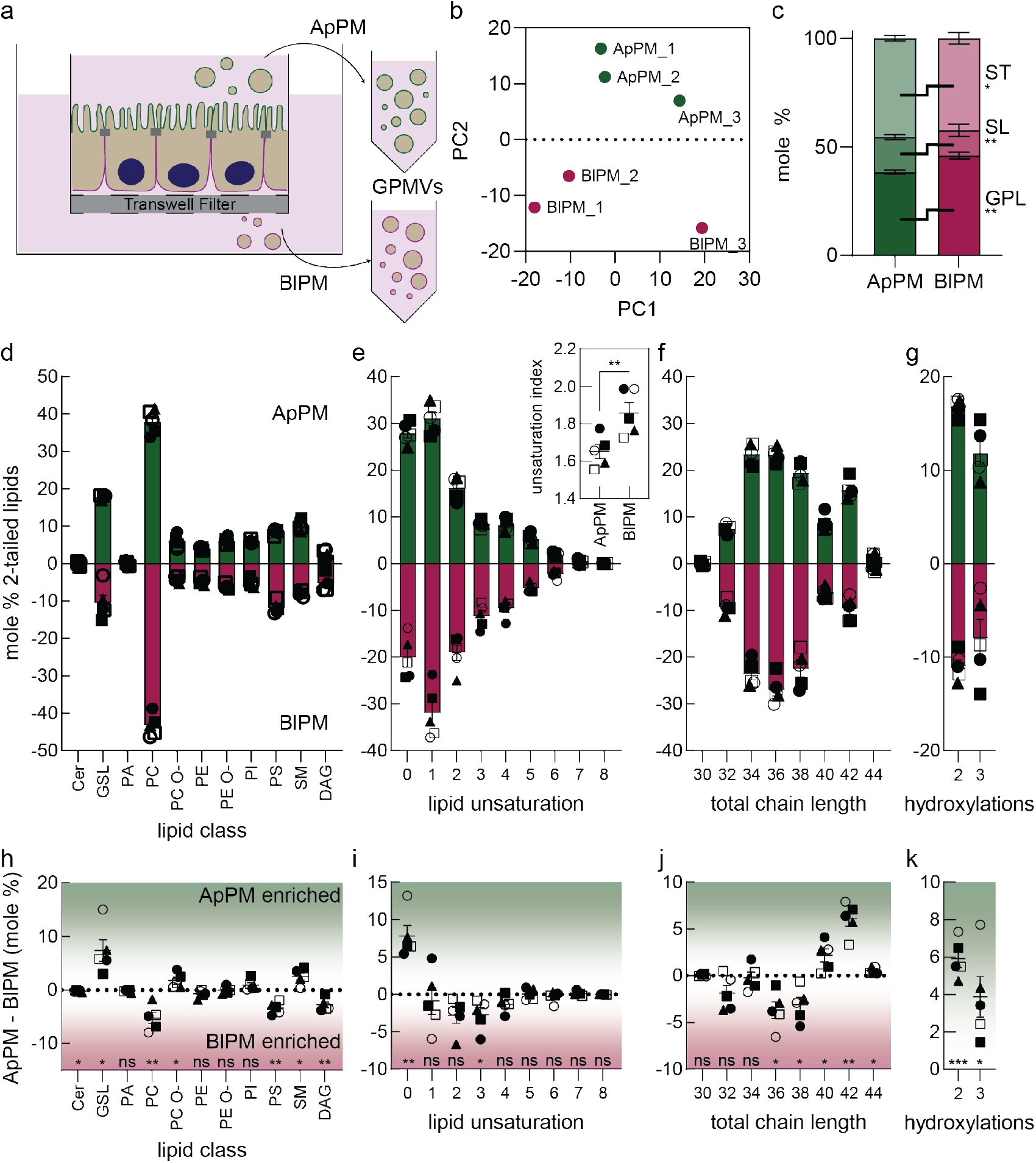
Apical and basolateral PMs have distinct lipidomes. (a) Schematic representation of GPMVs to separately isolate ApPM and BlPM lipidomes from either side of transwell filters. (b) Principal component analysis for lipidomics datasets from 13 days of polarization, symbol labels indicate paired samples. (c) Lipid structural categories in ApPMs and BlPMs. Shading indicates structural category: GPL - glycerophospholipid, SL - sphingolipid, ST - sterol. Paired t-test p values shown on right. (d-g) Phospholipid compositions (mol% of two-tailed lipids) by (d) headgroup, (e) acyl chain unsaturation, (f) chain length, and (g) hydroxylation. Phospholipid unsaturation index shown in inset of (e). (h-k) Differences between ApPM and BlPM lipidomes (d) headgroup, (e) acyl chain unsaturation, (f) chain length, and (g) hydroxylation. Paired samples are indicated with matching symbols, monolayers polarized for 13 days (n = 3, filled symbols) or 21 days (n = 2, open symbols). Mean and SEM shown, results of one sample t test shown in (e), ns – not significant, *p ≤ 0.05, **p ≤ 0.01, ***p ≤ 0.001. Abbreviations: ceramide (Cer), glycosphingolipid (GSL), sphingomyelin (SM), diacylglycerol (DAG), phosphatidic acid (PA), phosphatidylcholine (PC), phosphatidylethanolamine (PE), phosphatidylinositol (PI) and phosphatidylserine (PS), and the PE and PC ether derivatives (PE O-, PC O-).

Lipidomic analysis revealed substantial differences between ApPM and BlPM lipidomes. Principal component analysis revealed that ApPM and BlPM lipidomes are well separated, indicating that membrane polarity is a distinct and reproducible driver of lipidomic variance (Fig 1b). The ApPM was enriched in sphingolipids and cholesterol relative to the BlPM (SL and ST, respectively, Fig 1c). Correspondingly, the BlPM was enriched in glycerophospholipids relative to the ApPM (GPL, Fig 1c). The complete lipidomes of the ApPM and BlPM (Supporting Table 1) can be fractionated by lipid headgroup (Fig 1d), acyl chain unsaturation (Fig 1e), acyl chain length (Fig 1f), and acyl chain hydroxylation (Fig 1g). The ApPM had a significantly lower unsaturation index, i.e. the weighted average of lipid unsaturations for the lipidome (Fig 1e, inset), suggesting that its lipids were overall more saturated. We observed minimal lipidomic changes between 13 and 21 days of polarization (filled vs open symbols, Fig 1d-g), suggesting that compositions of fully polarized monolayers are stable over time. Generally, our observations are consistent with previous reports of lipidome remodeling during MDCK polarization^14^, as well as characterization of the apical PM lipidome isolated by physical ‘peeling’^13^.

The most striking differences between the ApPM and BlPM were in lipid features that have been previously associated with lipid rafts. Namely, cholesterol (ST, Fig 1c), glycosphingolipids (GSLs), and SM were significantly enriched in ApPM relative to the BlPM (Fig 1h). Conversely, the BlPM was enriched in PC and the signaling lipids PS and DAG relative to the ApPM. The enrichment of PS and DAG in the basolateral PM is consistent with their involvement in small G-protein and growth factor signaling^34,35^, which occur on the basolateral membrane surface. The ether-linked version of PC (PC O- or PC plasmalogen) was also enriched in the ApPM (Fig 1h), consistent with classical observations of plasmalogen enrichment in raft-associated detergent resistant membranes^36^. Further, saturated lipids (Fig 1i), those with very long (>40 carbons) acyl chains (Fig 1j), and hydroxylated lipids (Fig 1k) were significantly enriched in the ApPM relative to the BlPM. Altogether, our lipidomic measurements revealed that (a) apical and basolateral PM compartments have notably different lipidomes, (b) the ApPM is enriched in lipids that form tightly packed domains, and (c) that the BlPM is enriched in signaling lipids PS and DAG.

### Biophysical differences between PM compartments are consistent with polarized lipidomes

The differences between ApPM and BlPM lipidomes suggested that their membrane compartments may have distinct biophysical properties, with the apical being more tightly packed and ordered. To compare biophysical membrane phenotypes of the apical and basolateral PMs, we relied on lipophilic solvatochromic dyes, which have been extensively used to report on the biophysical properties of synthetic^37^ and natural membranes^38,39^. One example is Di4, whose fluorescence emission lifetime is highly sensitive to lipid packing^38,40–42^. Using fluorescence lifetime imaging microscopy (FLIM) to measure Di4 lifetime in live polarized MDCK monolayers, we found higher lifetimes in imaging planes containing apical surfaces compared to lateral and basal (Fig 2a,b), implying that the apical PM has tighter lipid packing than the basolateral PM. Using previously published calibrations, these differences in Di4 lifetimes correspond to ∼10% differences in area per lipid^41^.

**Figure 2.**
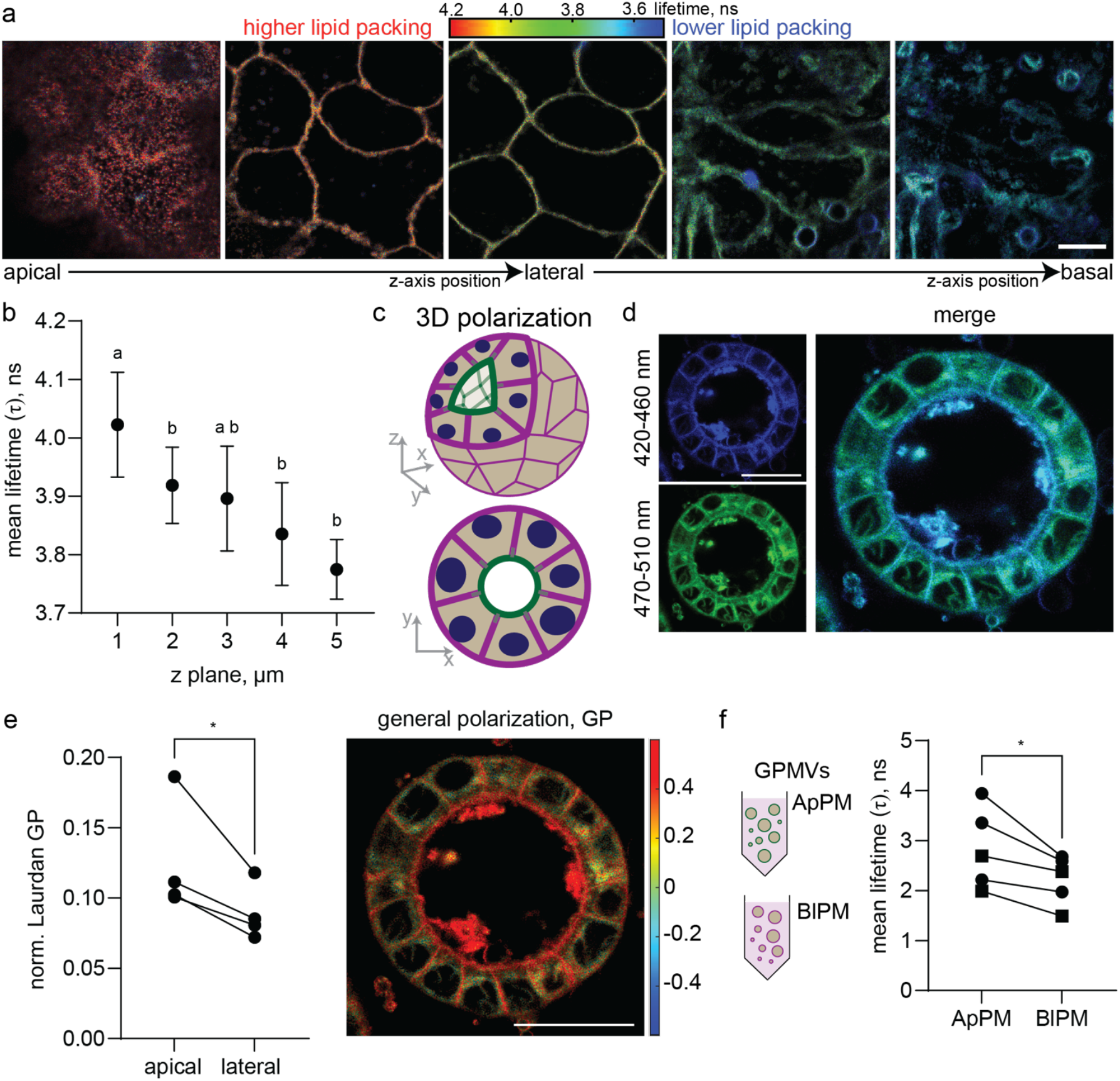
Polarized cell PM compartments differ in lipid packing. (a) Representative FLIM z-stack of a polarized MDCK monolayer stained with Di-4. (b) Quantified Di4 lifetime for various depths of the z-stack from apical (left) to basal (right) of polarized MDCKs monolayers as shown in (a); >3 fields of view per monolayer, n = 4 independent experiments, mean and SEM shown. Statistical comparisons were performed using one-way ANOVA with post-hoc multiple comparisons (Fisher’s LSD test); groupings are shown using the Prism Compact Letter Display, where groups that do not share a letter are significantly different (p<0.05). (c) Cartoon representation of cells grown as polarized spheroids. The apical surface (green) faces the lumen of the spheroid. (d) Representative confocal images of polarized spheroids stained with C-laurdan. (e) C-laurdan general polarization (GP) quantification of polarized spheroids. GP values of the PM are normalized by subtracting the GP of the inner cellular membranes (inner), >5 spheroids measured per experiment, mean of each experiment and paired t-test p value shown. Representative image shown on the right. (f) Di4 lifetimes for ApPM and BlPM GPMVs collected from monolayers polarized for 13 days (n = 2, squares) or 21 days (n = 3, circles). More than 10 GPMVs measured per replicate, p value for paired t-test shown. Scale bars in (a) is 5 µm and (d,e) are 25 µm.

To confirm this inference in a different experimental setting, MDCKs were grown in Matrigel, a three-dimensional extracellular matrix hydrogel that supports tissue mimetic morphogenesis. In Matrigel, individual MDCK cells proliferate and self-assemble into polarized spheroids with a cell-free lumen and clearly defined apical (lumen-facing) and basolateral (matrix-facing) PM compartments (Fig 2c and Supporting Fig 3a). In this setting, the apical surface is inaccessible to Di4, which cannot cross the PM or tight junctions (Supporting Fig 3b) due to its charged headgroup. Interestingly, we did detect small but significant differences in lipid packing between the lateral (i.e. cell-cell interfaces) and basal PM compartments (Supporting Fig 3b,c), with the basal PM being less packed than the lateral (as in the cells grown in two-dimensional cultures, Fig 2a,b). However, to compare these to the apical PM, it was necessary to switch to a different approach. Thus, we stained cells with another common solvatochromic membrane packing sensor, C-Laurdan, which is uncharged and therefore readily crosses membranes. C-Laurdan effectively stained both apical and basolateral PMs of the cells in spheroids (Fig 2d) and showed a blue-shifted emission spectrum, indicative of tighter lipid packing in the apical PM (Fig 2d, right). This effect can be quantified via the Generalized Polarization (GP), a dimensionless, normalized index of C-Laurdan’s red-shifted (490 nm) versus blue-shifted (440 nm) emission (Fig 2e, see Methods). The normalized GP of the lateral PM in MDCK spheroids was significantly lower than the apical, consistent with Di4 measurements in polarized MDCK monolayers. Finally, Di4 lifetime of GPMVs isolated from apical was greater than basolateral (Fig 2f), consistent with measurements of PMs in intact polarized cells (Fig 2a,b). Thus, across two different polarization contexts and two different probes, the apical PM showed tighter lipid packing than the basolateral PM. These biophysical differences are consistent with our lipidomics observations that apical PMs are rich in sphingolipids, cholesterol, long-chain saturated lipids, and hydroxylated lipids (Fig 1c,d-f,h-j), which together form tightly packed membrane domains^37,38^.

Finally, to directly assess whether lipid phase behavior is different in apical versus basolateral PMs, we again isolated ApPM or BlPM GPMVs and measured their miscibility transition temperature, which reports on their propensity to phase separate^32,43–45^. This measurement is performed by microscopically quantifying the fraction of vesicles that phase separate at various temperatures. GPMVs derived from both the apical and basolateral side of the monolayer (i.e. ApPM and BlPM, respectively) showed temperature-dependent liquid-liquid phase separation, indicating their capacity to form ordered and disordered membrane domains. The miscibility transition temperature of ApPMs was significantly lower than the BlPMs (Fig 3a,b), indicating significant differences in their propensity for phase separation. It is important to emphasize that this effect does not imply that there are ‘less rafts’ in these GPMVs or the apical PM. Rather, the lower miscibility transition temperature may result from reduced contrast in lipid domain order between coexisting membrane phases in the ApPM compared to the BlPM, a relationship supported by prior work showing that domain stability depends on the magnitude of the difference in lipid order between coexisting domains^33^.

**Figure 3.**
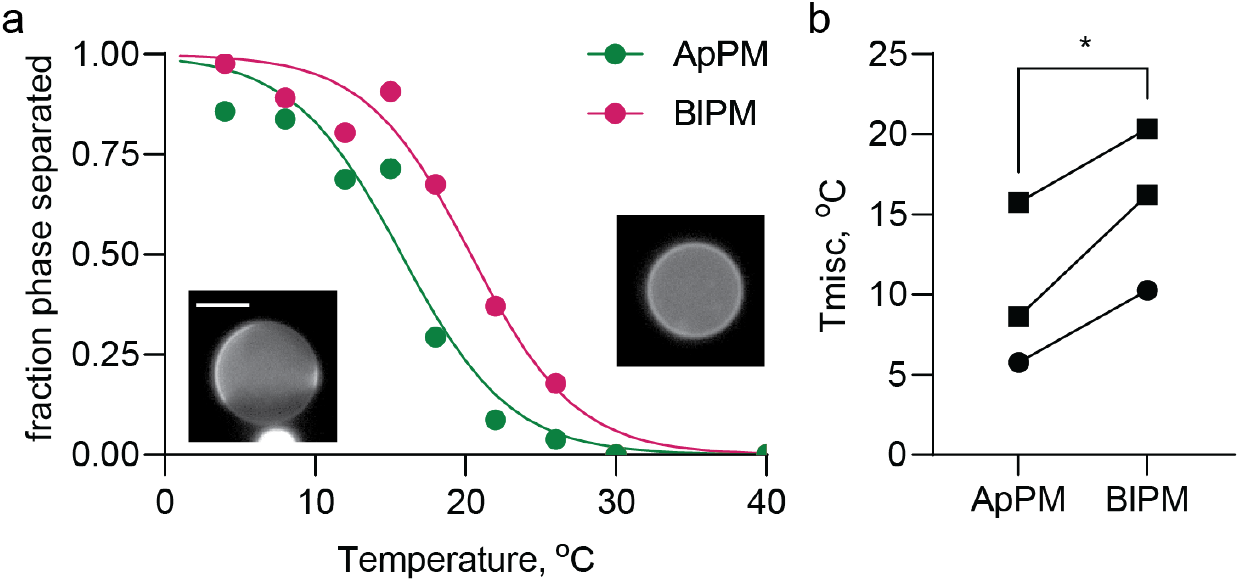
ApPM and BlPM have distinct phase separation. (a) Representative miscibility transition temperature data for ApPM and BlPM GPMVs and sigmoidal fit. (b) Miscibility transition (i.e. phase separation) temperature is slightly lower in ApPMs. GPMVs collected from monolayers polarized for 13 days (n = 2, squares) or 21 days (n = 1, circles). More than 10 GPMVs measured per temperature for each replicate, p value for paired t-test shown.

### Raft-preferring protein transmembrane domains are weakly enriched at the apical PM

To determine whether the differences in lipid composition and membrane biophysical properties between apical and basolateral PM compartments influence polarized protein sorting, we evaluated the steady-state localization of protein probes with known raft phase preferences. We define ‘raft phase preference’ by the partitioning of lipids and proteins into relatively ordered, cholesterol- and sphingolipid-rich phases of GPMVs^27,28^. To that end, GPMVs were observed under conditions of microscopic phase separation and stained with a lipophilic dye (Fast-DiO) that selectively partitions into the disordered/non-raft lipid phase. The phase preference of a protein of interest can then be quantified via its relative fluorescence intensity in the raft versus non-raft phase (Fig 4a), as previously demonstrated^31^.

**Figure 4.**
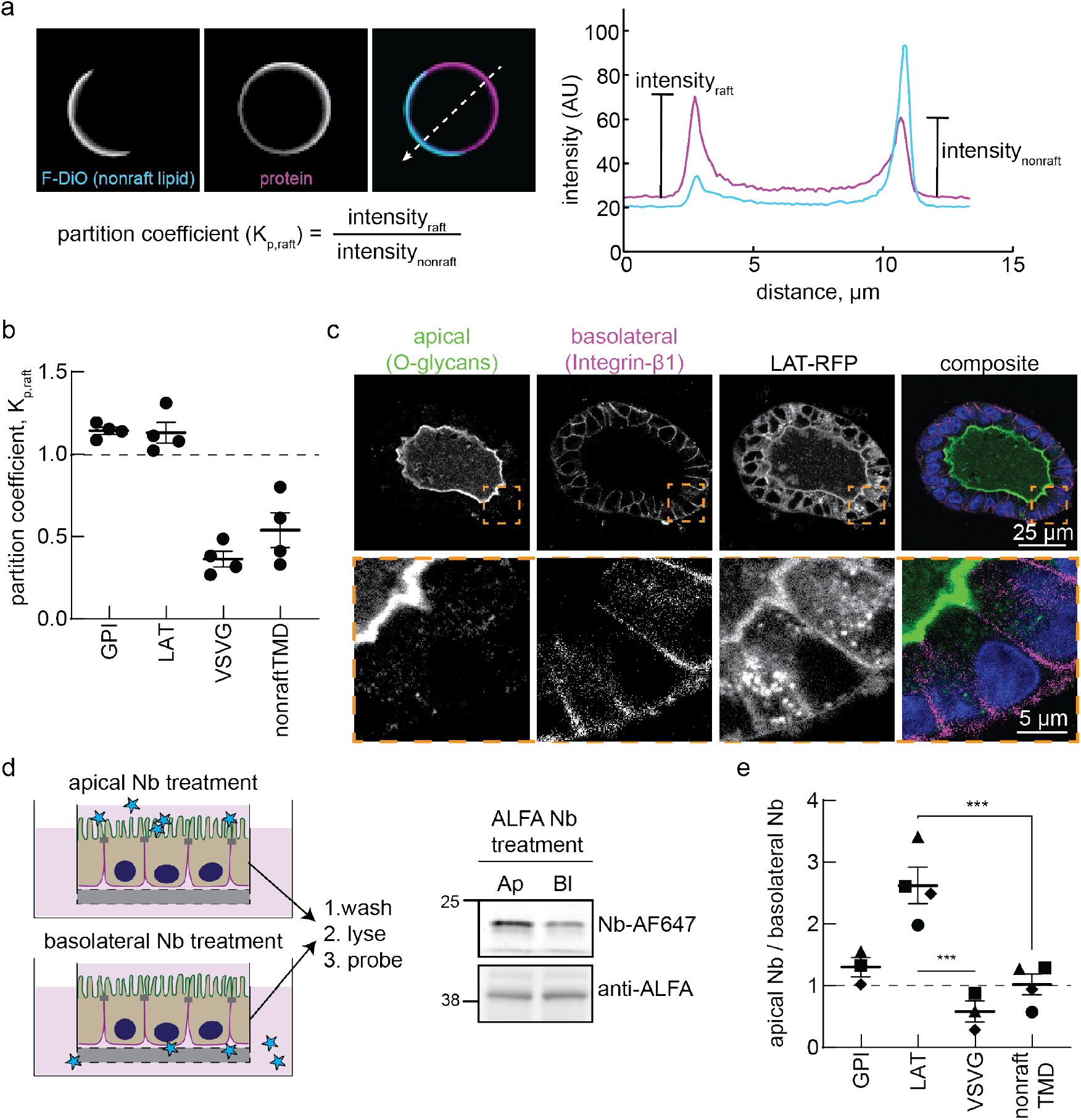
Raft-preferring proteins sort preferentially to the apical PM compartment. (a) Representative image of a phase separated GPMV, representative line scan for quantifying the relative intensity in the raft and non-raft region of the vesicle, and equation used to calculate partition coefficient (Kp). (b) Partition coefficients for four chimeric proteins of interest, each fused to a fluorescent tag. Mean and SEM shown, n = 4 experiments, > 10 GPMVs measured per experiment. (c) Representative confocal microscopy images of polarized MDCK ectopic expression cell line spheroids grown for 10+ days in Matrigel, fixed, and immunostained. Lower panels indicated zoomed in region of interest. (d) Schematic representation of biochemical labeling strategy to quantify relative apical and basolateral PM protein fractions. Briefly, polarized monolayers are stained from either the apical or basolateral media with fluorescent nanobodies to the ALFA tag. Unbound nanobody is washed off, monolayers are lysed, and lysates are probed for the amount of bound nanobody (Nb-AF647) and the amount of available ALFA tag (anti-ALFA). (e) Quantification of samples as shown in (d). Matching symbols indicate paired experiments, mean and SEM. Data were analyzed using one-way ANOVA with Fisher’s LSD post-hoc test for multiple comparisons. Individual p-values between groups are indicated.

We generated stable cell lines expressing various fluorescence-labeled proteins of interest, focusing on two natural, but not MDCK-native, single-pass transmembrane proteins (LAT and VSVG) that were expected to have opposite raft preference and two synthetic constructs: a minimal single-pass, transmembrane domain (TMD) construct that was engineered to have low raft affinity (nonraftTMD) and a GPI-anchored Halotag that has been extensively characterized as a raft marker. Quantification of raft partition coefficients (K_p,raft_) confirmed these predictions: both LAT and GPI had K_p,raft_ >1, revealing preference for more ordered phases, while VSVG and nonraftTMD were largely excluded from these domains (K_p,raft_ < 1; Fig 4b). To assess whether these differences in ordered phase affinity were related to polarized localization, we performed microscopy on MDCK spheroids expressing these protein probes using well-characterized apical and basolateral markers as spatial masks (Fig 4c). Qualitatively, we observed enrichment of LAT at the apical PM of polarized spheroids. However, for non-raft partitioning proteins, most signal was observed in intracellular compartments at steady-state (Supporting Fig 4a), consistent with previous findings that raft phase affinity facilitates PM localization in unpolarized cells^27,28^. Although there was observable signal at the PM with these constructs, this vesicular pool of non-raft proteins confounds the PM signal (Supporting Fig 4). Further, the apical PM is likely more convoluted than the basolateral, due to formation of microvilli. These issues inhibit meaningful, quantitative comparisons between apical and basolateral PM compartments by imaging.

We therefore devised a strategy for specifically labeling the PM fraction of a given protein of interest at either the apical or basolateral surface using extracellular ALFA tags. Here, the ALFA tag allows us to label only the external facing fraction of a construct by a fluorescent nanobody that cannot cross the PM. After labeling either the apical or basolateral side in live cells, cells were lysed and the fluorescent signal from the nanobody was quantified, thus revealing the relative abundance of the construct in the two PM compartments. These differences can then be normalized to the total expression of the tagged construct by western blotting the lysate against the ALFA tag (Fig 4d). This biochemical analysis revealed a significant enrichment of LAT in the apical PM (Fig 4d-e and Supporting Fig 5). The raft-preferring GPI-anchored-mCherry was also mildly apically enriched (Fig 4e)^1^. In contrast, neither of the two raft-excluded protein constructs (VSVG, nonraftTMD) were enriched in the apical PM but rather distributed approximately evenly between the two PM compartments. Thus, protein affinity for raft phases appears to impact preference for the relatively tightly packed apical PM compartment; however, it is important to emphasize that these differences were somewhat subtle, with both raft and non-raft proteins present in both PM compartments, suggesting that ordered domain affinity alone does not explain polarized protein localization.

**Figure 5.**
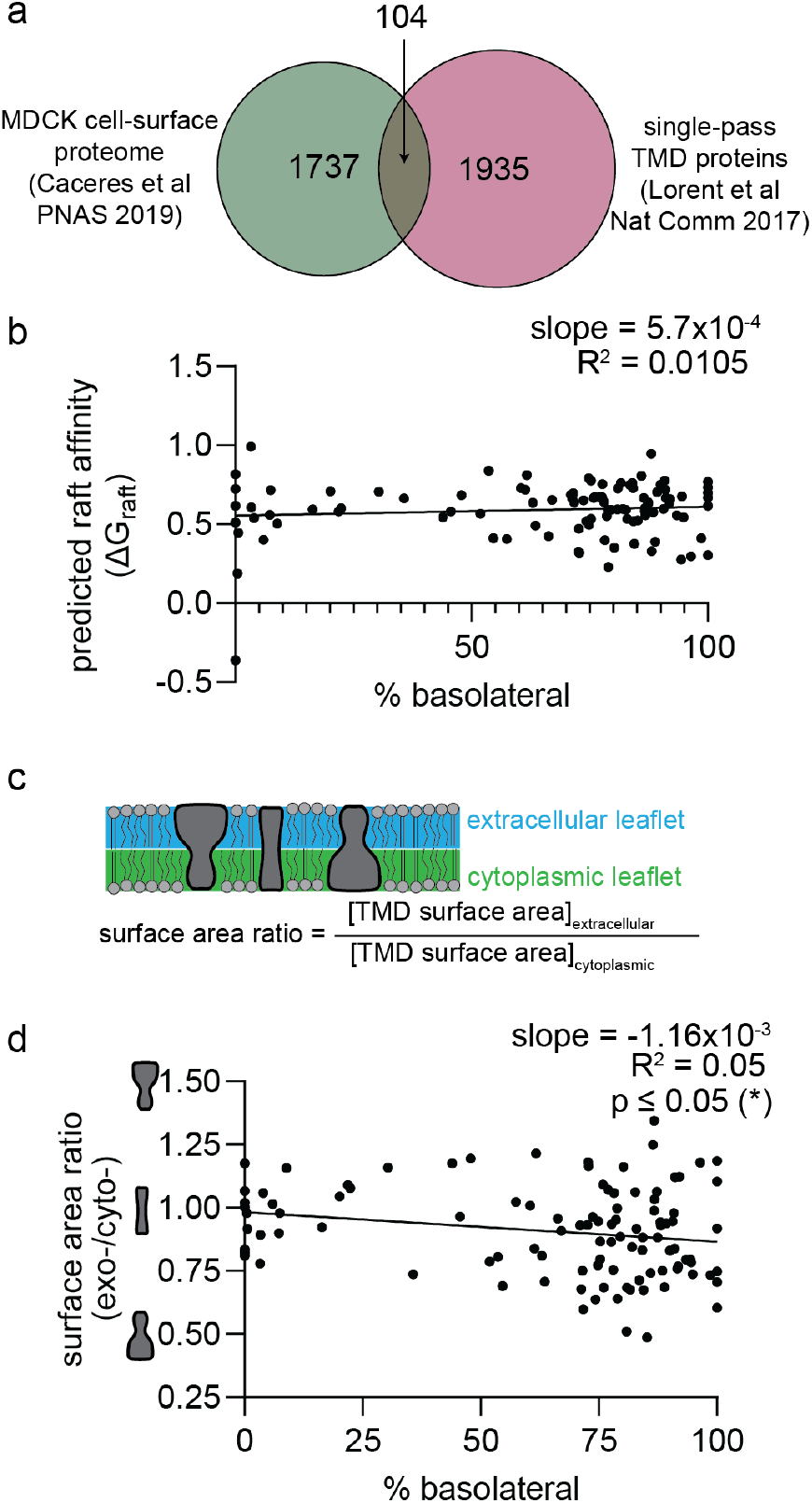
TMD features are minimally associated with polarized protein localization. (a) Venn diagram of proteins identified by Caceres et. al.^3^ via cell-surface biotinylation of polarized MDCK monolayers and proteins with predicted dG_raft_ (raft affinity)) from Lorent et. al.^28^ These two databases overlap for 104 unique, single-pass transmembrane proteins. (b) No significant correlation between dG_raft_ and polarized protein localization. (c) Schematic representation of transmembrane domain surface area asymmetry and calculated surface area ratio for transmembrane domains. (d) Correlation between TMD asymmetry and polarized protein localization for the 104 identified proteins has a weakly significant non-zero slope.

Despite the notable enrichment of raft-preferring TMD constructs at the apical PM (Fig 4c,e), we observed no substantial difference in the lipid-phase preference of the overall PM proteome between ApPM, BlPM, and naïve PM (Supporting Fig 6a-b). In this analysis, we non-specifically label all externally facing proteins and measure their raft phase preference in GPMVs, thereby capturing the ensemble behavior of the surface proteome rather than that of selected protein constructs. Non-specific labeling of external protein amine groups via NHS-AlexaFluor647 revealed similar raft partitioning for the aggregate surface proteome in apical, basolateral, and unpolarized GPMVs (Supporting Fig 6a). The values and trends were similar when external proteins were labeled using the lectin Wheat Germ Agglutinin (WGA), which binds glycosylated proteins (Supporting Fig 6b). Further, a representative single-pass transmembrane protein (LAT) had similar raft phase preference regardless of whether the partition coefficient is measured in ApPM, BlPM or naïve PM GPMVs (Supporting Fig 6c). These measurements indicate that GPMVs from the apical or basolateral surface can reveal protein preferences with lipid domains despite their different lipidomes.

Finally, comparing the predicted raft affinity from previously published work^28^ to the previously reported polarized cell proteome^3^, we find no correlation between raft affinity and polarized protein localization (Fig 5a-b and Supporting Table 2). Thus, preference for ordered lipid phases (as predicted by the model of Lorent et. al. ^28^) is not a major determinant of polarized membrane protein sorting. Interestingly, we observe a weak but significant correlation between TMD asymmetry and protein polarization (Fig 5c-d and Supporting Table 2), with more asymmetric TMDs (larger TMD surface area in the cytoplasmic leaflets) correlated with more basolateral localization.

## Discussion

In this study, we present a systematic comparison of the lipid composition, biophysical properties, and protein sorting between apical and basolateral PM compartments in polarized epithelial cells. We demonstrate that the apical PM is enriched in cholesterol, sphingolipids, and long-chain saturated lipids. This compositional profile corresponds to tighter lipid packing in the apical than the basolateral compartment. Moreover, we show that transmembrane proteins with high raft affinity exhibit mild preferential apical localization in polarized MDCK monolayers. These semi-synthetic probes allow us to probe raft-associated polarized protein sorting without confounding endogenous sorting signals. Together, these results suggest that the apical PM is enriched in raft-preferring lipids and proteins, supporting a model in which the collective, biophysical features of PM lipids contribute to the establishment and maintenance of epithelial membrane polarity. These data extend previous observations of the composition of the apical PM^12–14^ and provide, to our knowledge, the first direct comparisons between apical and basolateral PM compartments within the same polarized cellular assembly.

Importantly, both the lipidomics and biophysical approaches used here report on the bulk features of the PM compartments and do not provide direct evidence (or lack thereof) of nanoscale lateral heterogeneity, i.e. lipid rafts. One possible interpretation of our observations is that the apical PM contains a higher proportion of sub- microscopic, relatively ordered domains than the basolateral regions. This interpretation is potentially consistent with our observation that protein constructs with higher ordered domain affinity are somewhat enriched at the apical PM and previous work suggesting that the apical PM contains a continuous lipid raft phase^46^. However, our observations do not rule out that both compartments are laterally homogenous, with the apical being more tightly packed than the basolateral.

Despite the connections to raft-mediated lipid organization, our observations raise critical questions about the mechanisms of epithelial PM polarization and the maintenance of the apical PM. While the lipidomes of apical and basolateral compartments are clearly distinct, these differences are not consistent with a solely raft-based mechanism for apical PM biogenesis and sorting. Specifically, the apical PM is not fully ‘raft-like’ nor *vice versa* for the basolateral PM; the quantitative lipidomic differences between these compartments (Fig 1 and Supporting Table 1) are much more subtle than would be expected for differences between raft and non-raft phases in model membranes^29,45^. Similarly, while there is a relationship between raft affinity and apical PM preference for minimal protein probes, it is also relatively subtle (Fig 4e), with only ∼2-fold apical enrichment between the most and least raft-preferring probes. This mild enrichment is in contrast to the complete polarization observed for many PM proteins^3^. Similarly, we observe no meaningful relationship between predicted raft affinity of single-pass transmembrane proteins and their apical enrichment (Fig 5b). Thus, we conclude that raft-based sorting is unlikely to be the sole determinant of the distinctions between the apical and basolateral compositions of polarized MDCKs. Other possibilities for protein sorting determinants include cytoplasmic domain size^11^, glycosylation^47,48^, and other post-translational modifications^49^.

If raft-based sorting does not explain apical PM composition, why are tightly packing, raft-forming lipids enriched in the apical PM? One possibility is that the crucial barrier role of the apical PM in gut, kidney, and lung epithelial cells is mediated, at least in part, by its tightly packing lipidome. Sphingolipids, glycolipids, saturated lipids, and sterols pack tightly together, and this capacity may be supported by sphingolipid hydroxylation (Fig 1g,k). In this conception, enrichment of tightly packing lipids is an end in itself, rather than the means to sort other components.

Another important question is how apical enrichment of raft-associated lipids is generated. Clearly, certain proteins, including many GPI-anchored proteins^4,5,49^, are preferentially sorted to the apical PM. It is likely that these proteins have preferences for certain lipid components, even without invoking formation of stable, ordered, raft-like phases^18^. It is possible that highly specific mechanisms ensure the polarized localization of these proteins, leading to sorting of their associated lipids. An exciting example of this possibility is Mucin-1 (Muc1), the predominant glycoprotein of the glycocalyx in many polarized cells. Muc1 has high affinity for raft phases in GPMVs (Supporting Fig 6d), suggesting that it may recruit raft-preferring lipids during its trafficking to the PM.

We observed enrichment of DAG and PS lipids in the basolateral PM, which may reflect their role as second messengers for signaling processes occurring at the basolateral surface. Namely, DAG can be produced by the activity of phospholipase C and serve to activate protein kinase C^50^, both aspects being key nodes of signal transduction for growth factors and other extracellular signals^35^. PS has been implicated in electrostatic interactions with many peripheral proteins^51^, and in nanoclustering of small GTPases (e.g. Ras)^52–54^. In contrast, we observed an enrichment of sphingolipids and plasmalogens in the apical PM, consistent with previous observations^13,14^. Glycolipids likely play an important functional role in the apical PM glycocalyx and may direct apical protein traffic via lectin-mediated protein sorting mechanisms^55,56^. Plasmalogens have recently been implicated in membrane packing and curvature^57,58^, as well as resistance to oxidative damage^59,60^, both important functions for maintaining apical PM homeostasis. Future work could further explicate the role of membrane leaflet asymmetry in polarized cell function. A better understanding of membrane leaflet asymmetry could elucidate deeper distinctions between the apical and basolateral lipidomes, as several of the enriched/depleted species are likely to exist predominantly in only one leaflet (i.e., GSL in the outer leaflet or PS in the inner leaflet).

In conclusion, our findings establish links between membrane lipid composition, biophysical properties, and protein sorting in polarized epithelial cells. Quantitative comparisons of apical and basolateral membrane compartments suggest that lipid-mediated membrane organization may underlie aspects of polarized PM organization.

## Materials and Methods

### Key reagents

16% Paraformaldehyde Aqueous Solution, EM Grade (PFA, Electron Microscopy Sciences 69038-AS); 1,4-Dithiothreitol (DTT, Sigma-Aldrich 10708984001); Dulbecco’s Phosphate Buffered Saline, Modified (DPBS), w/o calcium (PBS, Sigma-Aldrich D8537-500ML); Fast DiO (Thermo Scientific D3898); Fast DiI (Thermo Scientific D7756); Goat serum (Thermo Scientific 31872); Prolong Diamond Antifade Mountant (Thermo Scientific P36961); Di4 (Di4 ANEPPDHQ, Thermo Scientific D36802); C-Laurdan (Tocris 7273); Licor Chameleon Duo Pre-stained Protein Ladder (LicorBio 928-60000); Precision Plus Protein Dual Color Standard (Bio-Rad 1610374EDU); PVDF membrane (Pierce 88520); Alexa Fluor 647 NHS Ester (Invitrogen A20006); Janelia Fluor 549 HaloTag Ligand (Promega GA1110); Halt Protease Inhibitor Cocktail (Thermo Scientific 78438); PMSF (Cell Signaling Technology 8553S); Matrigel Basement Membrane Matrix (Corning 354234)

### Antibodies/affinity reagents

anti-ZO1 Monoclonal (Thermo Scientific 33-9100); anti-Integrin Beta1 (EMD Millipore MABT409); PNA Lectin CF640R (Biotium 29063); anti-ATP1B1 (Sigma Aldrich SAB210573); anti-AE2 (GeneTex GTX1147); anti-Calnexin (Abcam ab22595); anti-rabbit IgG HRP linked (Cell Signaling Technology 7074S); Peroxidase (HRP) anti-mouse IgG (Cell Signaling Technology 7076S); Wheat Germ Agglutinin (WGA) CF640R (Biotium 29026); Goat anti-Mouse IgG (H+L) Cross-Adsorbed Secondary Antibody, Alexa Fluor 568 (Thermo Scientific A-11004); Goat anti-Rat IgG (H+L) Cross-Adsorbed Secondary Antibody, Alexa Fluor 488 (Thermo Scientific A-11006); ALFA antibody (Synaptic Systems N1580); ALFA sdAb FluoTag-X2 (Synaptic Systems N1502-AF647-L, RRID:AB_3075981); DAPI (NucBlue Live Ready Probes Reagent, Invitrogen R37605)

### Plasmids

HyperActive Transposase, Muc1-42TR-dCT-GFP pPB rtTA NeoR, pPB EF1a, pPB tetOn PuroR, pPB rtTA NeoR, and pLV Hygro TetOn were a gift from Matthew Paszek (Cornell University) and previously published^61^. To generate the pPB EF1a PuroR plasmid, the pPB EF1a plasmid was modified to include the puromycin resistance cassette and a custom multiple cloning site compatible with the existing library of Levental lab plasmids^28^. The puromycin resistance cassette including its promoter was PCR copied from pPB tetOn PuroR with 5’-TCGGCAATTCCCATGGAGCTGCAATAAACAAGTTGGGGTG-3’ and 5’-CGGAGCCGTCCCATGGTCCCCAGCAGGCAGAAGTATG-3’ and non-directionally inserted into the NcoI site of the pPB EF1a plasmid. The multiple cloning site was modified by annealing the following DNA duplex between the existing BamHI and XbaI restriction enzyme sites in the pPB EF1a plasmid: 5’-GGCAGGATCCGCTAGCCCCGGGACTAGTACCGGTAGATCTGAATTCGGCA-3’. Similarly, the pPB rtTA NeoR multiple cloning site was replaced to be compatible with existing restriction enzymes in the Levental lab. The pPB EF1 HygroR backbone was generated by PCR of the SV40 and Hygromycin resistance genes from pLV Hygro TetOn using 5’-TCGGCAATTCCCATGGTGTGTCAGTTAGGGTGTGG -3’ and 5’-CGGAGCCGTCCCATGCTATTCCTTTGCCCTCGGACG -3’ inserted non-directionally into the NcoI site of the pPB EF1a plasmid. LAT-TMD-RFP pPB EF1a PuroR and AllLTMD-RFP pPB EF1a PuroR (nonraftTMD) plasmids were generated by digesting the corresponding trLat and trAllL from Lorent et. al.^28^ with EcoRI and BamHI restriction enzymes and ligation into the pPB EF1a PuroR backbone. VSVG-RFP pPB EF1a PuroR was generated by PCR of the VSVG gene from Str-Golgin84_VSVG-SBP-EGFP (gift from Franck Perez; Addgene plasmid # 65305; http://n2t.net/addgene:65305; RRID:Addgene_65305)^62^ with 5’-ATCTACCGGTGAATTCATGAAGTGCCTTTTGTACTTAGCCT -3’ and 5’-AGGAGGCCATGGATCCCTTTCCAAGTCGGTTCATCTCT -3’, which was ligated into the LAT-TMD-RFP pPB EF1a PuroR plasmid between the EcoRI and BamHI restriction enzyme sites. To assemble GPI-Halo pPB EF1a HygroR, PCR the GPI-Halo fragment from GPI-Halo pcDNA3 with 5’-GTTGGGATCTACCGGTGCCACCATGGAGCTCTTTT -3’ and 5’-AAACGGGCCCTCTAGATTAAAGAACATTCATATACAGCACA -3’ and insert into the pPB EF1a HygroR plasmid between AgeI and XbaI restriction enzyme sites. ALFA exoplasmic tag plasmids through series of sequential steps. Frist, insert the AgeI to KpnI fragment of IL2ss-SBP-linker-LAT-TMD-RFP pcDNA3 from Castello-Serrano et. al.^63^ into the pPB EF1a PuroR plasmid. Swap the NheI to EcoRI fragment containing the SBP tag and linker for the following DNA duplex, which contains the ALFA tag sequence (underlined): 5’- CTAGCAGCCGTCTCGAGGAGGAGCTGAGAAGAAGACTGACCGAACCCGGGGGTGGTGGCAGTGGTGGTGGCAGTGGTGGTGGCAGTG -3’. Linkers were added to either side of the ALFA tag by digest by XmaI with non-directional insertion of the DNA duplex: 5’- CCGGGGGTGGTAGTGGTGGTGGCAGTGGTGGTGGCTCCGGTGGTAGTGGTGGTGGCAGTT -3’ and digestion by NheI with non-directional insertion of the DNA duplex: 5’- ctagTGGTGGTAGTGGTGGTGGCAGTGGTGGTGGCTCCGGTGGTAGTGGTGGTGGCAGTG -3’. Finally, the entire IL2ss-GGGSx5-ALFA-GGGSx8-LAT-TMD-RFP was swapped to the pPB rtTA NeoR backbone by restriction enzyme digest with AgeI and XbaI. AllL-TMD and VSVG-RFP versions were generated by swapping the EcoRI to BamHI fragments from the corresponding pPB EF1a PuroR plasmids to the IL2ss-GGGSx5-ALFA-GGGSx8-LAT-TMD-RFP pPB rtTA NeoR plasmid. Generate the IL2ss-GGGSx5-ALFA-GGGSx8-mCherry GPI pPB rtTA NeoR plasmid by PCR of the mCherry-GPI fragment from the mCherry-GPI pcDNA plasmid using 5’- TGGTGGCAGTGAATTCGTGAGCAAGGGCGAGGAGGATAA -3’ and 5’- AAACGGGCCCTCTAGATTAAAGAACATTCATATACAGCACA -3’ between the EcoRI and XbaI sites of the IL2ss-GGGSx5-ALFA-GGGSx8-LAT-TMD-RFP pPB rtTA NeoR plasmid.

Digested DNA fragments were joined using T4 DNA Ligase reactions (NEB M0202S). Restriction enzymes were purchased from NEB. PCR fragments were joined using In-Fusion (Takara Bio 638943). Plasmid sequences were confirmed by whole plasmid sequencing from Plasmidsaurus.

### Cell culture and manipulation

#### To culture cells

MDCK cells were obtained from ATCC. Cells were maintained in DMEM (Corning 10-013-CV) supplemented with 10% fetal bovine serum (Genesee Scientific 25-514H) and 1% penicillin-streptomycin (Sigma-Aldrich P4333-100 mL). Cells were maintained in a 37°C incubator with 5% CO_2_ and periodically tested for mycoplasma with the ATCC Universal Mycoplasma Detection Kit (30-1012K).

#### To generate ectopic expression cell lines

Low passage number MDCK cells were co-transfected by electroporation (Mirus Bio MIR 50117 electroporation solution) with the corresponding gene of interest plasmid and the HyperActive Transposase plasmid (provided by Matthew Paszek, Cornell University)^61^. 24-48 hours post transfection, cellular medium was changed to include 1.5 µg/mL puromycin (Sigma Aldrich P8833) for 10 days or 750 µg/mL G418 (Thermo Scientific 10131035) for 14 days before withdrawing the selection reagent.

#### To generate polarized monolayers

Low passage number MDCK cells or derivative, ectopic-expression cell lines of MDCK parental cells were plated onto 24 mm transwell filters with 3 µm pores (Corning 3414) with 200,000 cells per transwell filter. Media was changed every other day on both sides of the transwell filter. Cells were grown for the indicated number of days before experiments.

#### To generate polarized spheroids

Low passage number MDCK cells or derivative, ectopic-expression cell lines of MDCK parental cells were suspended as a single-cell suspension in 5% Matrigel containing-medium and overlayed onto a glass bottom dish (Matsunami Glass D141400) pre-coated with 50 µL of polymerized Matrigel. Medium containing 5% Matrigel was substituted every other day and cells were grown for 10 or more days for all experiments.

#### Lipidomics sample collection and analysis

For GPMV lipidomics samples, mature monolayers were transferred to new, sterile 6 well plates to remove any cells which had migrated to the bottom of the original 6 well plate. GPMV solution was prepared with 0.072% PFA, 4 mM DTT, 150 mM NaCl, 10 mM Hepes, 2 mM CaCl_2_, pH 7.4. Three washes were performed with 2 mL of GPMV solution on each side of transwell filters. Transwells were incubated with GPMV solution for 45 minutes at 37°C. GPMVs were collected from each side of the transwell filter. Apical GPMV samples were filtered with Ultrafree Centrifugal Filters (EMD Millipore UFC30SV00) to remove cell debris. GPMV samples were pelleted at 16,000 rcf for 2 h at 4°C. The supernatant was aspirated, and the pellet was resuspended in 250 uL of PBS. Samples were stored at -72°C until shipped on dry ice to Lipotype (Dresden, Germany). Lipid features include an internally controlled standard used for pmol quantification of lipid species, except for Forssman glycolipid (FGL). Per Gerl et. al.^13^ and Sampaio et. al.^14^, FGL is assumed to be present at 8 mol% for ApPM lipidomes polarized for 13 or more days. Using this assumption and based on the mass spectroscopy peak intensities for FGL lipid species, the FGL pmol is calculated for the ApPM samples. The ratiometric relationship between the ApPM FGL intensity and the calculated ApPM FGL pmol is used to determine the FGL pmol for the corresponding BlPM sample based on its measured FGL intensity. These calculated FGL pmol are incorporated into the mole % two-tailed lipids (corrected) values. To obtain #db (corrected), one double bond which exists in the backbone structure of the lipid species, is removed from the total number of double bonds measured for SL.

#### Lipid nomenclature

The following lipid names and abbreviations are used: ceramide (Cer), glycosphingolipid (GSL), sphingomyelin (SM), diacylglycerol (DAG), lactosyl ceramide (DiHexCer), glucosyl/galactosyl ceramide (HexCer), sterol ester (SE), Forrsman glycolipid (FGL) and triacylglycerol (TAG), as well as phosphatidic acid (PA), phosphatidylcholine (PC), phosphatidylethanolamine (PE), phosphatidylglycerol (PG), phosphatidylinositol (PI) and phosphatidylserine (PS), as well as their respective lysospecies (lysoPA, lysoPC, lysoPE, lysoPI and lysoPS) and ether derivatives (PC O-, PEp, LPCp and LPEp). Lipid species were annotated according to their molecular composition as follows: (lipid class)-(sum of carbon atoms in the fatty acid side chains):(sum of double bonds in the fatty acid side chains);(sum of hydroxyl groups in the long-chain base and the fatty acid moiety), for example, SM-32:2;1. Where available, the individual fatty acid composition according to the same rule is given in brackets (for example, 18:1;0-24:2;0).

#### PCA analysis

Principle component analysis was performed on complete lipidomics data from ApPM and BlPM isolated on day 13 (the values for each specie were calculated as mol% of total lipids measured). The data was analyzed using the prcomp function in RStudio with scaling of the data.

#### Immunostaining and imaging

After polarization of monolayers, using a blade, transwell filters were cut away from the transwell insert. Monolayers were fixed and processed on their filter fragments. Polarized spheroids were fixed and processed in their culture chamber. Samples were washed with PBS, fixed with 4% PFA for 15-20 minutes at room temperature, and rinsed again with PBS. Samples were blocked with 5% goat serum in PBS with 0.3% Triton X-100 for 1 hour at room temperature. Samples were incubated with the indicated primary antibodies or lectin diluted 1:200 in 5% goat serum, PBS, 0.3% Triton-X overnight at 4°C. Samples were subsequently washed and incubated with NucBlue and corresponding fluorescent secondary antibodies diluted 1:500 in 5% goat serum, PBS, 0.3% Triton-X for 2 hours at room temperature. Samples were washed and monolayers are mounted overnight onto a number 1.5 coverslip using Prolong Diamond. Samples were then imaged on a Leica SP8 confocal using a 63x water objective (NA 1.20).

#### Flow Cytometry of GPMVs

Cells were grown in transwell filters for 13 days or 6 well cell culture plates for less than 24 hours (naïve). Cells were stained with Fast-DiO or Fast-DiI at 50 ug/mL in 150 mM NaCl, 10 mM Hepes, 2 mM CaCl_2_, pH 7.4 for 8 minutes at 4°C. Fast-DiO or Fast-DiI were selectively applied to one side of the transwell filter only. Subsequently, GPMVs were formed and collected from each sample as described above. Fluorescence of GPMV samples was measured on a BD LSRFortessa. Compensation was applied to the data, and data was analyzed in FCS Express.

### Western blotting

*For unpolarized cells lysates:* Samples were plated and lysed within 24 hours using RIPA lysis buffer with protease inhibitor cocktail and PMSF with cell scraping.

#### For GPMV samples

Sample GPMVs were generated as described from either polarized monolayers or naïve cells and pelleted at 16,000 rcf for 2 h at 4°C. The supernatant was aspirated, and the pellet was resuspended in 250 uL of using RIPA lysis buffer with protease inhibitor cocktail and PMSF.

#### For polarized protein localization lysates

Cell lines ectopically expressing the indicated proteins of interest with extracellular ALFA tags were polarized on transwell filters for 13 days as described. Before lysis, 4 µg/mL doxycycline (Sigma Aldrich D9891) was added to the media on either side of the transwell filter to induce expression of the protein of interest. After 18 hours of incubation, transwell samples were removed from the incubator and washed three times with complete media and transferred to a new 6 well plate to eliminate potential contamination from any cells that had landed on the bottom of the original plate. Samples were incubated either from the top or bottom side of the transwell filter with 2 mL of media containing 15 nM Nb-AF647 (ALFA sdAb FluoTag-X2) for 20 minutes at 4°C. Nanobody labeling was performed at a concentration previously confirmed to be saturating based on flow cytometry experiments; varying Nb concentrations had no effect, suggesting that Nb saturated the available ALFA binding sites. Transwells were subsequently washed with PBS and lysed using RIPA lysis buffer with protease inhibitor cocktail and PMSF with cell scraping.

Lysed samples were incubated on ice for at least 15 minutes. Then, samples were sonicated using a probe tip sonicator for three cycles of 10 seconds on, 10 seconds off at 30-40% amplitude and stored at -72°C until used. BCA was performed on cell lysates (Thermo Scientific 23223). Note that the DTT in GPMV samples is present at an unknown final concentration and confounds quantification by BCA assay, therefore GPMV samples were loaded at a constant volume. Samples were run in BioRad precast gels and transferred by Trans-Blot Turbo to PVDF membrane. Membranes were blocked with 3% BSA in TBST for more than 30 minutes at room temperature. Primary antibodies were incubated on membranes at manufacturer recommended dilutions overnight at 4°C, and secondary antibodies were incubated on membranes for more than two hours at room temperature. Membranes were imaged using Clarity ECL Substrate (Bio-Rad 1705061) on Azure Biosystems 300Q imager. Fluorescent antibodies were imaged using Licor Odyssey Infrared Imager before blocking step.

### Fluorescence Lifetime Imaging Microscopy (FLIM) with Di4

#### Live monolayer imaging

After polarization of monolayers, using a blade, transwell filters are cut away from the transwell insert. Monolayers were rinsed with GPMV buffer (150 mM NaCl, 10 mM Hepes, 2 mM CaCl_2_, pH 7.4) and incubated with 1 ug/mL Di4 diluted in GPMV buffer for 8 minutes at room temperature. Stained monolayer filter fragments were imaged within 20 minutes at room temperature with a Leica SP8 confocal microscope with a 63x water immersion objective (NA 1.20). A 488 nm 80 MHz pulsed white light laser excitation was used and photons were collected above 550 nm. Z-stacks were acquired with a 1 µm step size. Plasma membrane regions of interest were fit using Leica’s proprietary software (LASX) for a two-component n-exponential reconvolution, and the intensity weighted mean lifetime value is reported for the region of interest for each z-plane.

#### GPMV imaging

GPMVs were isolated from polarized monolayers as described and allowed to settle at room temperature. The bottom fraction of the sample was isolated and stained with a 1:1000 dilution of Di4 stock. After 20 minutes, the bottom fraction of the sample was transferred to a BSA coated coverslip and imaged according to the FLIM settings described above.

#### Live spheroid imaging

After polarization of spheroids, samples were washed three time with GPMV buffer (150 mM NaCl, 10 mM Hepes, 2 mM CaCl_2_, pH 7.4) and incubated with 1 µg/mL Di4 in GPMV buffer for 45 minutes at 37°C. Samples were subsequently imaged as describer within 20 minutes.

### Microscopic quantification

#### Of protein of interest localization by confocal

A custom macro was developed in Fiji (ImageJ) to automate the processing and quantification of confocal microscopy images. For each image series, the macro optionally applies background subtraction using Otsu’s auto-thresholding method to generate a binary mask, which is then multiplied with the original image to isolate the signal of interest. The resulting image is converted to 32-bit, and all background pixels (value = 0) are replaced with NaN to exclude them from downstream measurements. The macro then selects the full image area and records intensity measurements. All steps, including file loading, masking, background removal, thresholding, and data export, are batch-processed across multiple user-specified image series to ensure consistent and reproducible analysis.

#### Of miscibility transition temperature

Miscibility transition temperatures (Tmisc) were quantified as previously described^31,64,65^. Briefly, phase separation was microscopically assessed for >10 vesicles/temperature and the fraction of phase-separated vesicles plotted against temperature was fit with a sigmoidal curve.

#### Of Generalized Polarization with C-Laurdan

After polarization of spheroids, samples were fixed with 4% PFA as described and subsequently stained with 5 µM C-Laurdan in GPMV buffer (150 mM NaCl, 10 mM Hepes, 2 mM CaCl_2_, pH 7.4) for 3 hours at room temperature. Samples were washed three times for 5 minutes each with GPMV buffer before being imaged at room temperature with a Leica SP8 confocal microscope with a 63x water immersion objective (NA 1.20). Samples were illuminated with a 405 nm Diode laser and fluorescent signal was collected from 420 – 460 nm and 470 – 510 nm. MATLAB (The MathWorks, Natick, MA) was used to calculate the two-dimensional (2D) GP map, where GP for each pixel was calculated from a ratio of the two fluorescence channels as previously described^15^. Briefly, each image was binned (2 × 2), background subtracted, and thresholded to keep only pixels with intensities greater than 3 standard deviations of the background value in both channels. The GP image was calculated for each pixel using the equation below.

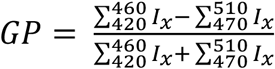

#### Of raft partition coefficients

Partition coefficient measurements were performed as previously described^28,31,66^. Briefly, live cell membranes were stained with 5 μg/ml of Fast-DiO, a green fluorescent lipid dye that strongly partitions to disordered phases^67^. Following staining, GPMVs were isolated from cells as described. To quantify protein partitioning, GPMVs were observed on an inverted epifluorescence microscope (Leica) below 10°C. The partition coefficient (K_p,raft_) for each protein construct was calculated from fluorescence intensity of the fluorescent construct in the raft (I_Lo_) and non-raft (I_Ld_) phase.

## Statistical analysis

Statistical tests were performed as indicated in figure legends using GraphPad Prism software.

## Data and code availability

All data and code used for analysis will be provided upon request.

## Acknowledgements

We thank Barbara Diaz-Rohrer and Allison Skinkle for their help with initial protocol development for polarized GPMV extraction and Professor Matthew Paszek (Cornell University) for mucin and piggybac plasmids. Funding for this work was provided by the NIH/National Institute of General Medical Sciences (GM134949 and AI183581 to I.L. and F32GM145028 to C.R.S.). The flow cytometry data for this manuscript were generated in the University of Virginia Flow Cytometry Core Facility (RRid:SCR_017829) and is partially supported by the NCI Grant (P30-CA044579).

## Author Contributions

C.R.S., K.R.L., and I.L. designed research, analyzed data, and wrote the manuscript.

C.R.S. and K.R.L. performed experiments.

