## Supplementary Figures and Table Legends for "Apical and basolateral plasma membranes in epithelial cells have distinct lipidomes and biophysical properties"

File contains 6 Supplementary Figures and associated legends; also, two Legends for Supplementary Tables

### Table Legends

**Supplementary Table I: Lipidomes of apical and basolateral GPMVs (ApPM and BlPM, respectively).** Complete membrane lipidomes from individual samples are shown at two different times for polarization (13 days and 21 days).

**Supplementary Table II: Comparison between polarized (apical versus basolateral) localization and TMD features.** The table includes information for ~100 transmembrane proteins whose apical versus basolateral localization was quantified by Caceres et al and whose raft affinity was predicted by Lorent et al based on the structures of their single-pass transmembrane domains.

### Supplementary Figures and Legends

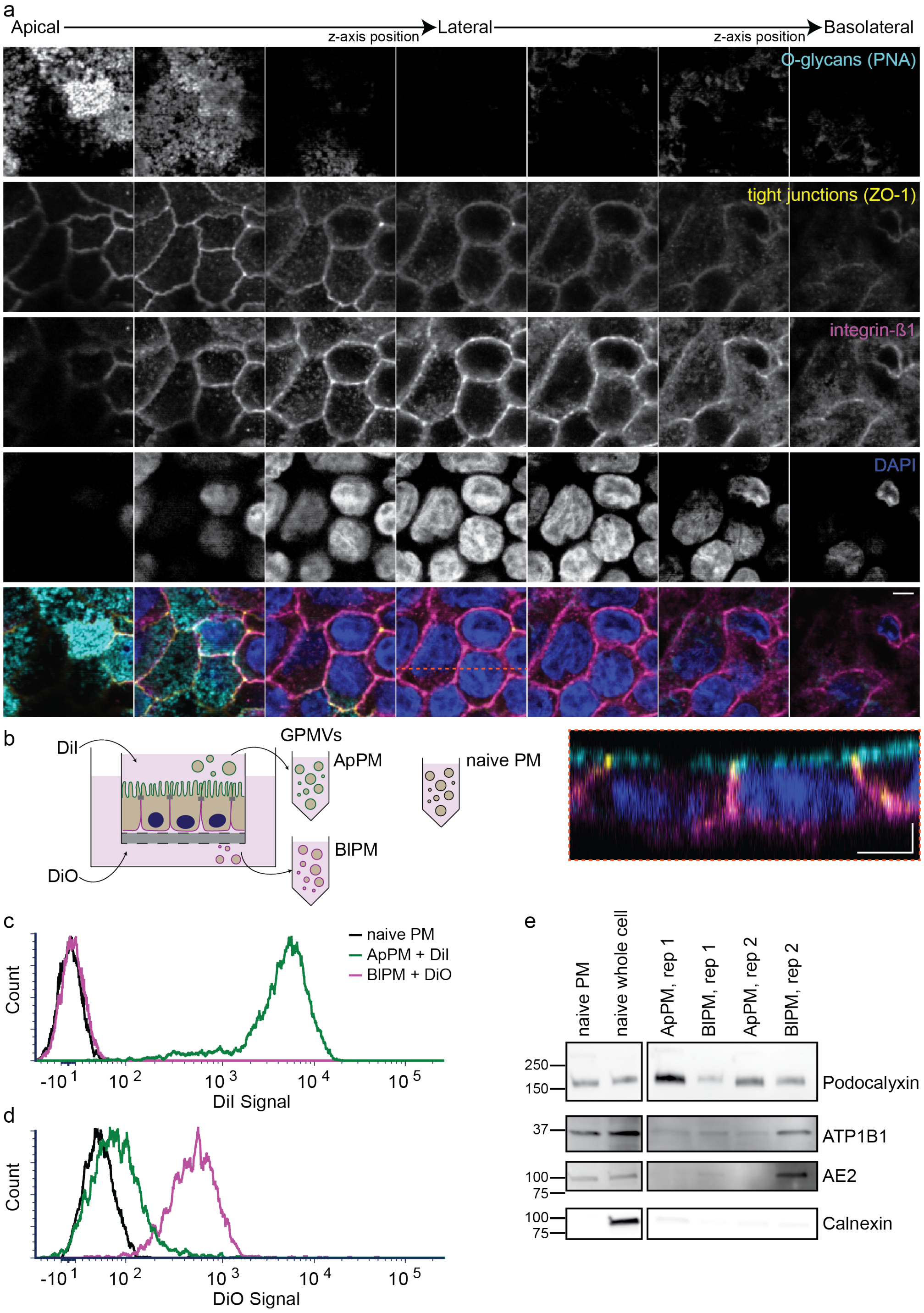


**Supporting Figure 1: Validation of PM sample preparation from polarized monolayers.** (a) Representative confocal images of polarized MDCK cultures grown on transwell filters for 13 days. Horizontal scale bars are 4 µm, vertical is 2 µm. (b) Schematic of lipophilic dye staining onto either side of transwell filter polarized monolayers to test lipid mixing between apical and basolateral PM compartments during GPMV isolation. (c) Representative flow cytometry histogram for DiI signal: while the apical GPMVs are brightly stained with DiI, basolateral GPMVs have no DiI signal above background, suggesting no lipid mixing. (d) Representative flow cytometry histogram for DiO signal. Inversely, bright DiO staining is only observed in the basolateral but not the apical GPMVs. These observations reveal minimal lipid mixing between apical and basolateral compartments during GPMV isolation. (e) Western blot validation of ApPM and BlPM. ApPM samples are enriched in the apical PM resident protein, podocalyxin. BlPM samples are enriched in the basolateral PM resident proteins, ATP1B1 and AE2. All PM samples are depleted of the ER resident Calnexin.

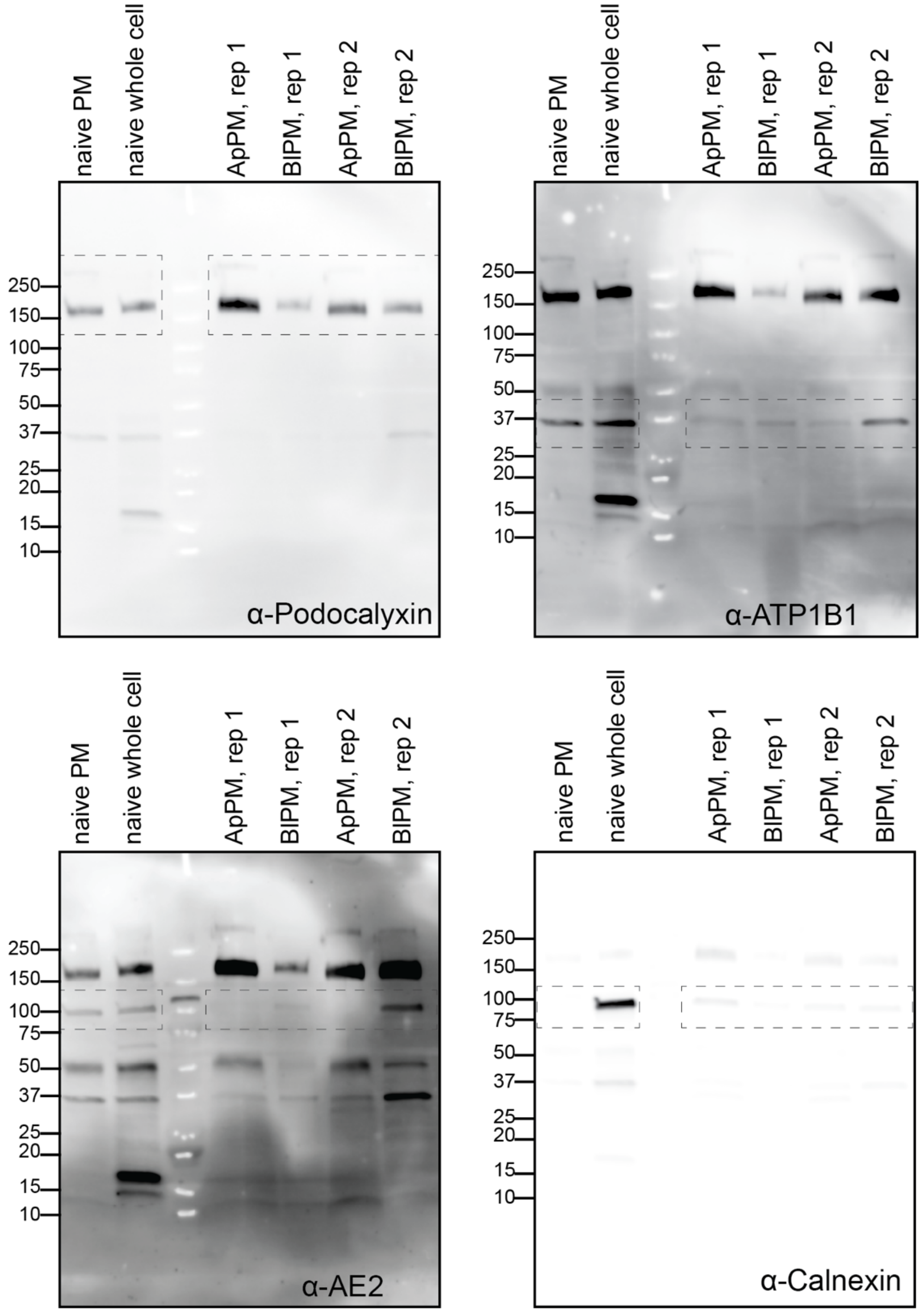


**Supporting Figure 2: Complete Western blot images associated with Supporting Figure 1.** Complete Western blot images associated with cropped ROIs shown in Supporting Figure 1e. The same membrane is probed sequentially, 1 – podocalyxin, 2 – ATP1B1, 3 – AE2, and 4 – calnexin. The ladder is run in the third lane.

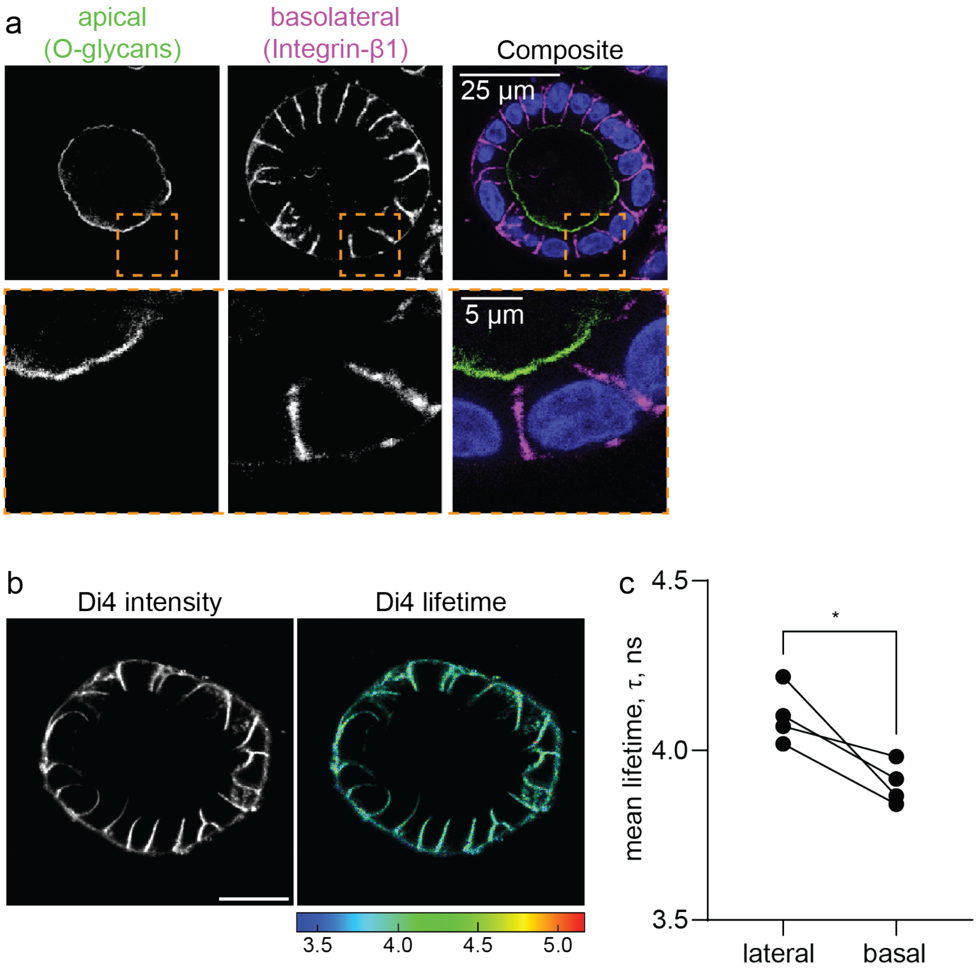


**Supporting Figure 3: Quantification of PM packing in polarized spheroids by Di4.** (a) Representative immunofluorescence of apical (peanut agglutinin, i.e. O-glycans, green) and basolateral (integrin-β1, magenta) markers and cellular nuclei (blue). Bottom row is zoom-in region from top row. (b) Representative image of outer leaflet Di-4 staining on spheroids showing lateral and basal staining while dye is excluded from the apical membrane compartment. (c) Quantification of Di-4 lifetime shows slightly higher lipid packing in lateral versus basal PMs of spheroids. >5 spheroids per sample, paired t-test p value shown. Scale bar in (b) is 20 µm.

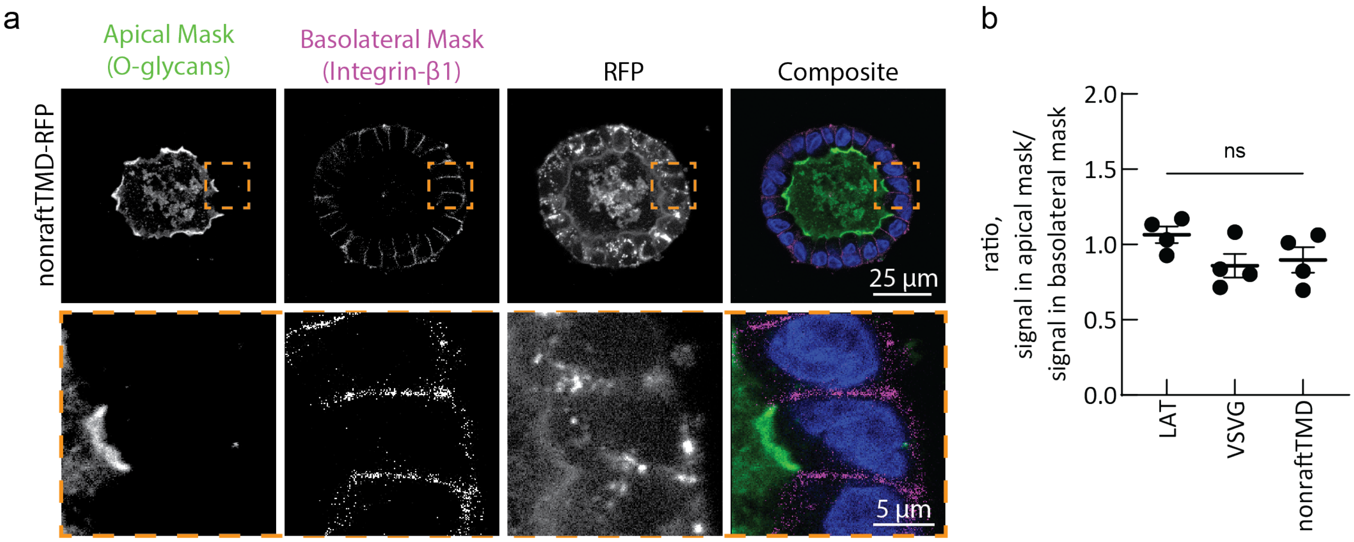


**Supporting Figure 4: Non-raft TMD localization by confocal microscopy.** (a) Representative confocal microscopy images of polarized MDCKs (grown for 10+ days in Matrigel) in spheroids ectopically expressing nonraftTMD-RFP (immunostaining for polarization markers). Lower panels indicated zoomed in region of interest. (b) Quantification of RFP signal from apical or basolateral mask regions in microscopy images, > 4 spheroids per experiment, n = 4 experiments, mean and SEM shown. One-way ANOVA with Fisher's LSD post-hoc test for multiple comparisons reveals no significant differences between the constructs.

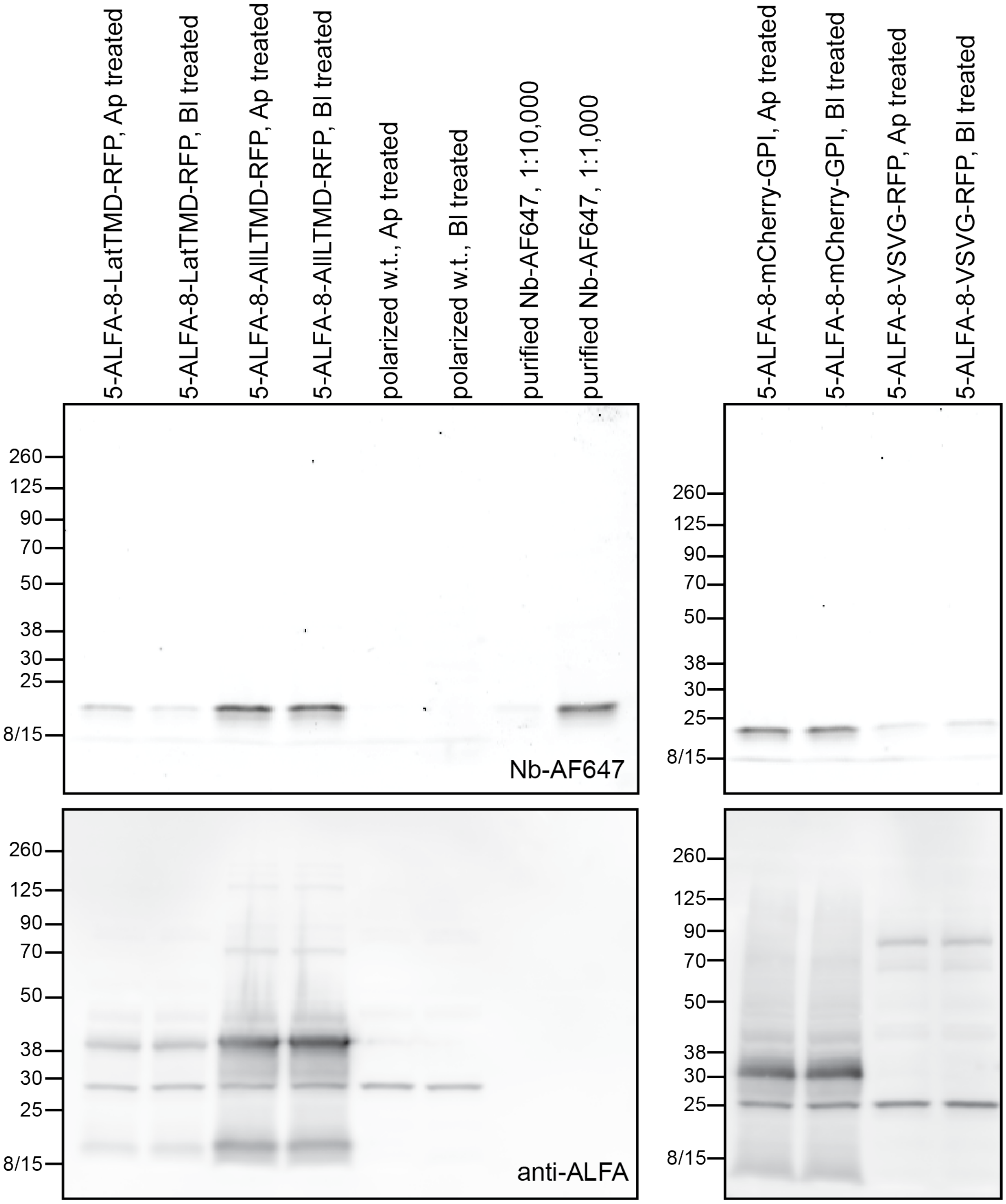


**Supporting Figure 5: Complete fluorescence and Western blot results.** Shows complete, representative PVDF membrane fluorescence for bound ALFA Nanobody-Alexa Fluor 647 (Nb-AF647, top row) and Rabbit-anti-ALFA tag Western blot (anti-ALFA, bottom row). The Nb-ALFA appears at the expected size of ~18 kD and is consistent with the band observed for the purified Nb-AF647 at both dilutions. MDCK cells that do not express ALFA-tagged constructs do not bind the Nb-AF647, as expected and indicated by the absence of the 18 kD band (top left image). These cells also lack the ALFA-binding epitope and show only a non-specific band in the anti-ALFA blot (bottom left image), which is present across all cell lysate samples. Expected protein of interest molecular weights: 5-ALFA-8-LatTMD-RFP and 5-ALFA-8-AllLTMD-RFP ~ 37 kD; 5-ALFA-8-mCherry-GPI ~ 39 kD; and 5-ALFA-8-VSVG-RFP ~ 89 kD.

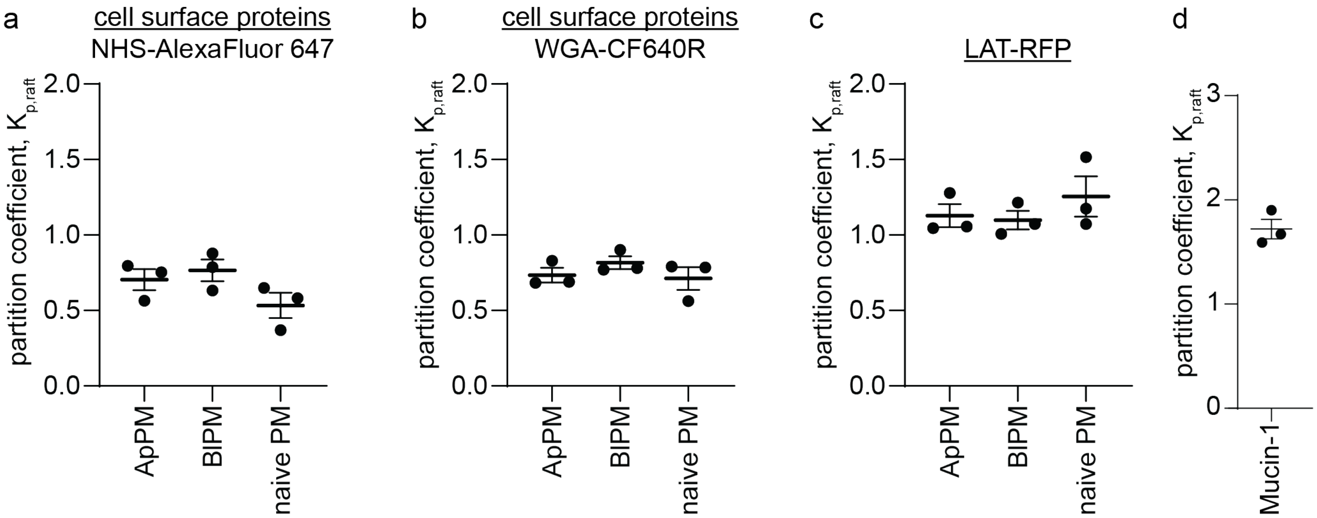


**Supporting Figure 6: Partition coefficients in GPMVs from various cellular compartments.** (a) Overall partition coefficient ($K_{p,raft}$) for PM proteins stained non-specifically with an amine-reactive dye (NHS-AlexaFluor 647) in ApPM, BlPM, or naive PM samples. (b) Partition coefficient ($K_{p,raft}$) for proteins stained broadly by wheat germ agglutinin (WGA-CF640R) in ApPM, BlPM, or naive PM samples. (c) Partition coefficient for LAT-TMD-RFP in ApPM, BlPM, or naive GPMVs. Mean and SEM are shown for three independent biological replicates with >4 GPMVs per replicate. (d) $K_{p,raft}$ for the abundant, apically localized glycoprotein Mucin-1.
